# DNA-guided establishment of canonical nucleosome patterns in a eukaryotic genome

**DOI:** 10.1101/013250

**Authors:** Leslie Y. Beh, Noam Kaplan, Manuel M. Müller, Tom W. Muir, Laura F. Landweber

## Abstract

A conserved hallmark of eukaryotic chromatin architecture is the distinctive array of well-positioned nucleosomes downstream of transcription start sites (TSS). Recent studies indicate that *trans*-acting factors establish this stereotypical array. Here, we present the first genome-wide *in vitro* and *in vivo* nucleosome maps for the ciliate *Tetrahymena thermophila*. In contrast with previous studies in yeast, we find that the stereotypical nucleosome array is preserved in the *in vitro* reconstituted map, which is governed only by the DNA sequence preferences of nucleosomes. Remarkably, this average *in vitro* pattern arises from the presence of subsets of nucleosomes, rather than the whole array, in individual *Tetrahymena* genes. Variation in GC content contributes to the positioning of these sequence-directed nucleosomes, and affects codon usage and amino acid composition in genes. We propose that these ‘seed’ nucleosomes may aid the AT-rich *Tetrahymena* genome – which is intrinsically unfavorable for nucleosome formation – in establishing nucleosome arrays *in vivo* in concert with *trans*-acting factors, while minimizing changes to the coding sequences they are embedded within.

## Introduction

Nucleosomes are the fundamental packaging unit of eukaryotic chromatin. Each nucleosome consists of ~147bp of DNA wrapped around a histone octamer (Luger et al. 1997). Nucleosomes serve as versatile signaling platforms, through the installation and removal of histone post-translational modifications (Zentner and Henikoff 2013) and histone variants (Maze et al. 2014). The organization of nucleosomes across the genome also plays an important regulatory role as it lowers the physical accessibility of DNA to cellular factors. For example, increasing evidence indicates that nucleosome positioning within functional elements across the genome directly impacts DNA-based transactions, such as transcription (Lam et al. 2008; Piña et al. 1990). In light of this, it is crucial to understand how nucleosomes are organized across the genome.

The rapid development of high-throughput sequencing technologies has permitted genome-scale surveys of nucleosome organization in every major eukaryotic model system, including *C. elegans*, *Drosophila*, humans, zebrafish, and both budding and fission yeast. These maps revealed strikingly similar nucleosome patterns near gene starts, where a nucleosome-depleted region upstream of the TSS is followed by a stereotypical array of nucleosomes inside the gene (Yuan et al. 2005; Lee et al. 2007; Mavrich et al. 2008b; Chang et al. 2012; Lantermann et al. 2010; Chen et al. 2013b; Zhang et al. 2014). Recent studies have used the budding yeast *Saccharomyces cerevisiae* as a model to understand nucleosome positioning mechanisms underlying the stereotypical nucleosome pattern near eukaryotic TSSs. In principle, nucleosome organization can be guided both by the intrinsic DNA sequence preferences of histone octamers, and by the action of *trans*-acting factors (Struhl and Segal 2013; Zhang et al. 2011; Hughes et al. 2012; Kaplan et al. 2009; Zhang et al. 2009). These two mechanisms, which are distinct, but not mutually exclusive, have been studied by comparing nucleosome positions across the genome *in vivo* and *in vitro*. The *in vitro* nucleosome maps were generated by reconstituting nucleosomes on yeast genomic DNA, in the presence or absence of *trans*-acting factors, represented by cell extracts or ATP-dependent chromatin remodelers (Kaplan et al. 2009; Zhang et al. 2011, 2009). Such experiments revealed that *trans*-acting factors, rather than the DNA sequence preferences of nucleosomes (Gkikopoulos et al. 2011; Yen et al. 2012; Zhang et al. 2009, 2011; Hughes et al. 2012), mainly underlie the characteristic nucleosome array downstream of TSSs. This stands in contrast to the nucleosome-depleted regions upstream of TSSs, which were found to be intrinsically unfavorable to nucleosome formation (Kaplan et al. 2009; Zhang et al. 2009). These findings have since been considered the consensus in the field (Struhl and Segal 2013).

Here, we dissect the respective contributions of nucleosome sequence preferences and *trans*-acting factors to nucleosome organization in the somatic macronuclear genome of the model ciliate *Tetrahymena thermophila*. The *Tetrahymena* genome is very GC-poor (22% GC), second only among eukaryotic genomes to *Plasmodium falciparum* (Gardner et al. 2002). It exhibits an unconventional structural organization, with ~225 unique chromosomes, each amplified to ~45n (Eisen et al. 2006). *Tetrahymena* is a widely established model for understanding chromatin biology, having provided seminal contributions to current knowledge of histone variants and post-translational modifications (Brownell et al. 1996; Allis et al. 1980). Recent work continues to reveal novel connections between chromatin modifications and diverse biological processes in *Tetrahymena*, ranging from DNA elimination (Liu et al. 2007) and replication (Gao et al. 2013), to the maintenance of genome integrity (Papazyan et al. 2014). However, no genome-scale analysis of chromatin has been reported in *Tetrahymena* to date, with data on nucleosome positioning being restricted to only a few loci (Cech and Karrer 1980; Palen and Cech 1984). In order to characterize nucleosome organization and dissect their underlying positioning mechanisms in *Tetrahymena*, we performed genome-wide MNase-based nucleosome mapping on log-phase and starved cells, as well as on histones assembled on sheared naked *Tetrahymena* DNA *in vitro* (Supplemental Figs. S1 and S2). These data together represent, to our knowledge, the first global analysis of chromatin structure in a ciliate.

In contrast to previous studies, we unexpectedly observe the stereotypical *in vivo* pattern in the *in vitro* nucleosome map of *Tetrahymena*. Another surprising finding arose through the systematic analysis of individual genes, where we discover that only subsets of the nucleosomes are positioned in individual genes *in vitro*. We find that *in vivo* such nucleosomes are flanked by well-positioned nucleosomes. Additionally, these sites coincide with locally GC-rich DNA, which is intrinsically favorable for nucleosome formation. Importantly, these constraints exert biases on codon usage and amino acid composition, because the DNA-encoded nucleosomes usually reside within the coding regions of genes. In light of these data, we propose a mechanism in which DNA-guided nucleosomes act as seeds to aid the establishment of *in vivo* nucleosome arrays in genes, while minimizing the impact on overlapping coding sequences.

## Results

### Genome-wide nucleosome maps of the *Tetrahymena thermophila* macronuclear genome

We established comprehensive maps of nucleosome organization in the *Tetrahymena* macronuclear genome through MNase-seq across two different nutritional conditions *in vivo*. In addition, we performed MNase-seq on reconstituted *Tetrahymena* chromatin *in vitro*, obtained by assembling histone octamers on sheared naked genomic DNA, in the absence of any other *trans*-acting factors (see Methods and Supplemental Fig. S1). Direct comparisons of the *in vivo* and *in vitro* datasets allow the inference of distinct nucleosome positioning mechanisms acting on the genome. For all analyses, we analyzed nucleosome positioning, rather than nucleosome occupancy, by assessing the distribution of nucleosome dyads across the genome, as inferred from the mid-points of individual nucleosome sequencing reads (see Methods).

First, we verified that MNase-seq datasets exhibit high coverage of the *Tetrahymena* nucleosome landscape by subsampling reads at varying proportions, and subsequent nucleosome calling using DANPOS (Chen et al. 2013a). The detected number of nucleosomes approached saturation well before full sampling of each MNase-seq dataset, indicating that *Tetrahymena* nucleosomes are well-sampled (Supplemental Fig. S2). We measured the nucleosome repeat length as 199bp, agreeing well with previous estimates obtained from gel analysis of MNase-treated macronuclear chromatin (Gorovsky et al. 1978, and Supplemental Fig. S3A). The average nucleosome linker length remained constant in both log-phase and starved nutritional conditions. This differs from previous studies suggesting that organisms exhibit an evolutionarily conserved increase in nucleosome spacing in response to starvation (Chang et al. 2012).

We then validated our MNase-seq datasets by analyzing nucleosome positions at the 5’ non-transcribed spacer of the *Tetrahymena* ribosomal DNA (rDNA) locus. Well-positioned nucleosomes flank both origins of replication within the 5’ NTS *in vivo* (Supplemental Fig. S4), closely corroborating independent studies that mapped nucleosomes at this locus through Southern analysis (Palen and Cech 1984). We observe similar patterns of nucleosome positioning in both log-phase and starved chromatin, consistent with previous reports (Palen and Cech 1984). Interestingly, our data suggest that the proximal origin of replication is not nucleosome-free, and is instead occupied by a nucleosome that is susceptible to increased MNase digestion at elevated temperatures (Supplemental Fig. S4). No evidence of nucleosome positioning could be detected *in vitro*, indicating that the distinctive chromatin organization of the rDNA locus arises from *trans*-acting factors, possibly associated with replication machinery.

### *Tetrahymena* exhibits stereotypical nucleosome patterns near TSSs *in vivo*

Eukaryotic nucleosome organization is most distinct near the 5’ ends of genes, where regularly spaced nucleosomes lie downstream of a nucleosome depleted region (Yuan et al. 2005; Lee et al. 2007; Mavrich et al. 2008b; Chang et al. 2012; Lantermann et al. 2010; Chen et al. 2013b). We find that this pattern is conserved in both *Tetrahymena* and yeast, with TSSs lying in the GC-poor nucleosome depleted region (Fig. 1 and Supplemental Fig. S5). The pattern in *Tetrahymena* is maintained between different nutritional conditions *in vivo* (Supplemental Fig. S5). We then investigated the relationship between transcription and nucleosome organization. Gene expression correlates with higher nucleosome density (defined by the average number of nucleosome centers per unit length of DNA) downstream of the TSS (Supplemental Fig. S6) in log-phase cells. By contrast, it correlates with lower nucleosome density ~250bp upstream of the TSS (Supplemental Fig. S7). Both trends are more apparent in the log-phase rather than the starved nutritional condition. These results suggest that processive transcription contributes to nucleosome density downstream of TSSs, consistent with previous studies in yeast (Hughes et al. 2012). On the other hand, an open chromatin environment upstream of TSSs could facilitate the binding of core transcriptional machinery to promoter elements for subsequent transcription. However, the contribution of such effects to nucleosome organization may vary according to the environmental context.

**Figure 1.**
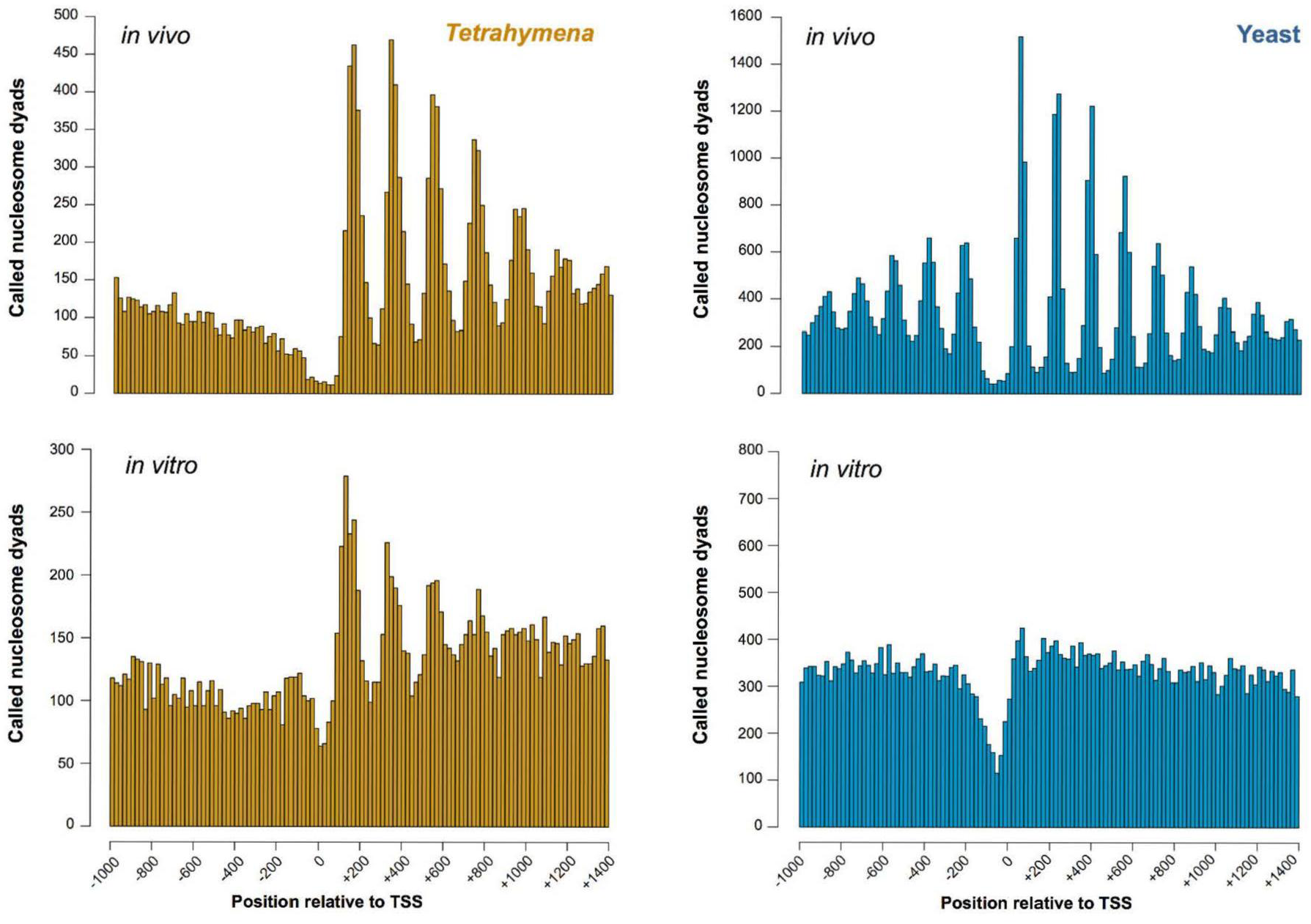
*in vivo*-like nucleosome organization without *trans*-acting factors. Histograms of nucleosome positions relative to the TSS were computed from yeast and *Tetrahymena* MNase-seq data using the same bioinformatic pipeline. A phased distribution of nucleosome positions downstream of the TSS is observed in chromatin from *Tetrahymena* and yeast grown in rich media. Surprisingly, an *in vivo*-like pattern of nucleosome positioning is observed *in vitro* for *Tetrahymena*, but not yeast.

### The stereotypical nucleosome array is present *in vitro* in *Tetrahymena* but not in yeast

We then compared *in vitro* organization around the TSS, between *Tetrahymena* and yeast. Unlike the *in vivo* data, we surprisingly find that the *in vitro* nucleosome patterns were markedly different between *Tetrahymena* and yeast (Fig. 1). Reconstituted *Tetrahymena* nucleosomes preferentially occupied positions that closely resemble the *in vivo* pattern (Fig. 1, Supplemental Fig. S5). We also observed that *in vitro* nucleosome peaks were less distinct and slightly shifted upstream relative to their matching *in vivo* peaks. No such *in vitro* organization was observed in yeast.

Following this, we performed several controls to validate this unusual observation. In order to rule out the possibility that the observed nucleosome organization *in vitro* arose from over-amplification during PCR, we removed duplicate reads from MNase-seq datasets and analyzed the distribution of nucleosome dyads around TSSs. The distinct organization of *in vitro* nucleosome dyads persisted even when duplicate reads were removed from MNase-seq datasets, ruling out this possibility (Supplemental Fig. S8). Our finding is also robust over a wide range of parameters used for nucleosome calling (Supplemental Fig. S9). Sequencing of MNase-digested naked *Tetrahymena* DNA did not show such a pattern (Supplemental Fig. S5), confirming that the nucleosome pattern observed *in vitro* does not result from biases in MNase cleavage.

Using these data, we found that nucleosome positioning is more similar between *in vivo* and *in vitro* datasets near TSSs, compared to other locations in the genome (Supplemental Fig. S10), reinforcing the notion that endogenous DNA sequences play an especially important role in organizing chromatin within *Tetrahymena* genes.

### Individual genes possess subsets of nucleosomes *in vitro*, rather than the complete array

It is important to realize that aggregate analysis of genomic data can be misleading. Specifically, the fact that we recover a well-positioned nucleosome array after averaging over many genes does not neccesarily imply that such an array exists in individual genes. We thus asked whether the unexpected similarity between average nucleosome patterns *in vivo* and *in vitro* also holds at the level of individual genes in *Tetrahymena*. To address this, we systematically measured the prevalence of positioned +1, +2, and +3 nucleosomes across the genome (Table S1; see Methods) by analyzing nucleosome patterns in individual genes. We henceforth term these nucleosomes as ‘canonical nucleosomes’. Strikingly, we find that most genes possess a subset of these canonical nucleosomes *in vitro*, rather than completely recapitulating the average pattern. A large fraction of genes (71.6 %) exhibit at least one canonical nucleosome *in vitro*, only slightly lower than that *in vivo* (76.3 %). In contrast, 32.5 % of genes had two canonical nucleosomes *in vitro*, compared to twice as many genes *in vivo* (61.7 %). Only a minority of genes (8.9 %) had nucleosomes at all three canonical positions *in vitro*, while this was much more extensively observed *in vivo* (43.8 %). Additionally, phasogram analysis did not show evidence of regular nucleosome arrays *in vitro* (Supplemental Fig. S11). Thus, unexpectedly, the average *in vivo*-like pattern that we observe *in vitro* is mainly explained by nucleosomes occupying various subsets of canonical positions within individual genes, rather than all positions near the TSS. This is clearly observed in profiles of nucleosome organization within individual genes (Supplemental Fig. S12).

### GC-rich sequences underlie DNA-guided nucleosomes in *Tetrahymena* genes

Since *in vitro* nucleosomes were reconstituted in the absence of *trans*-acting factors, we asked what DNA sequence preferences of nucleosomes underlie their stereotypical distribution near TSSs *in vitro*. GC content has previously been identified as a major component of such sequence preferences (Tillo and Hughes 2009). In particular, AT-rich sequences, such as poly(dA: dT) tracts, are refractory to nucleosome formation (Field et al. 2008; Nelson et al.; Suter et al. 2000; Segal and Widom 2009). Similar to other eukaryotes, we observe a decrease in GC content at TSSs, coinciding with nucleosome-depleted regions (NDRs) *in vitro* and *in vivo* (Fig. 2). Due to the low histone: DNA concentration used for reconstitution (4:10), the size of sheared DNA used in reconstitution (0.85-2 kb), and our observation that subsets of canonical nucleosome positions are occupied *in vitro*, we conjectured that local sequence features specifically located downstream of the TSS could underlie nucleosome organization *in vitro*, rather than previously suggested statistical concentration-based nucleosome positioning effects (Kornberg and Stryer 1988; Mavrich et al. 2008a). We thus examined the nucleotide composition of individual genes whose *in vitro* nucleosome maps show *in vivo*-like nucleosome organization. Notably, these genes exhibit oscillations in GC content downstream of the TSS, with an average amplitude of 1 % – 2 % GC and a period of ~200bp, coincident with canonical nucleosome positions (Fig. 2). The data collectively suggest that GC content oscillations within *Tetrahymena* coding sequences may contribute to regularly spaced nucleosomes *in vitro* and *invivo*.

**Figure 2.**
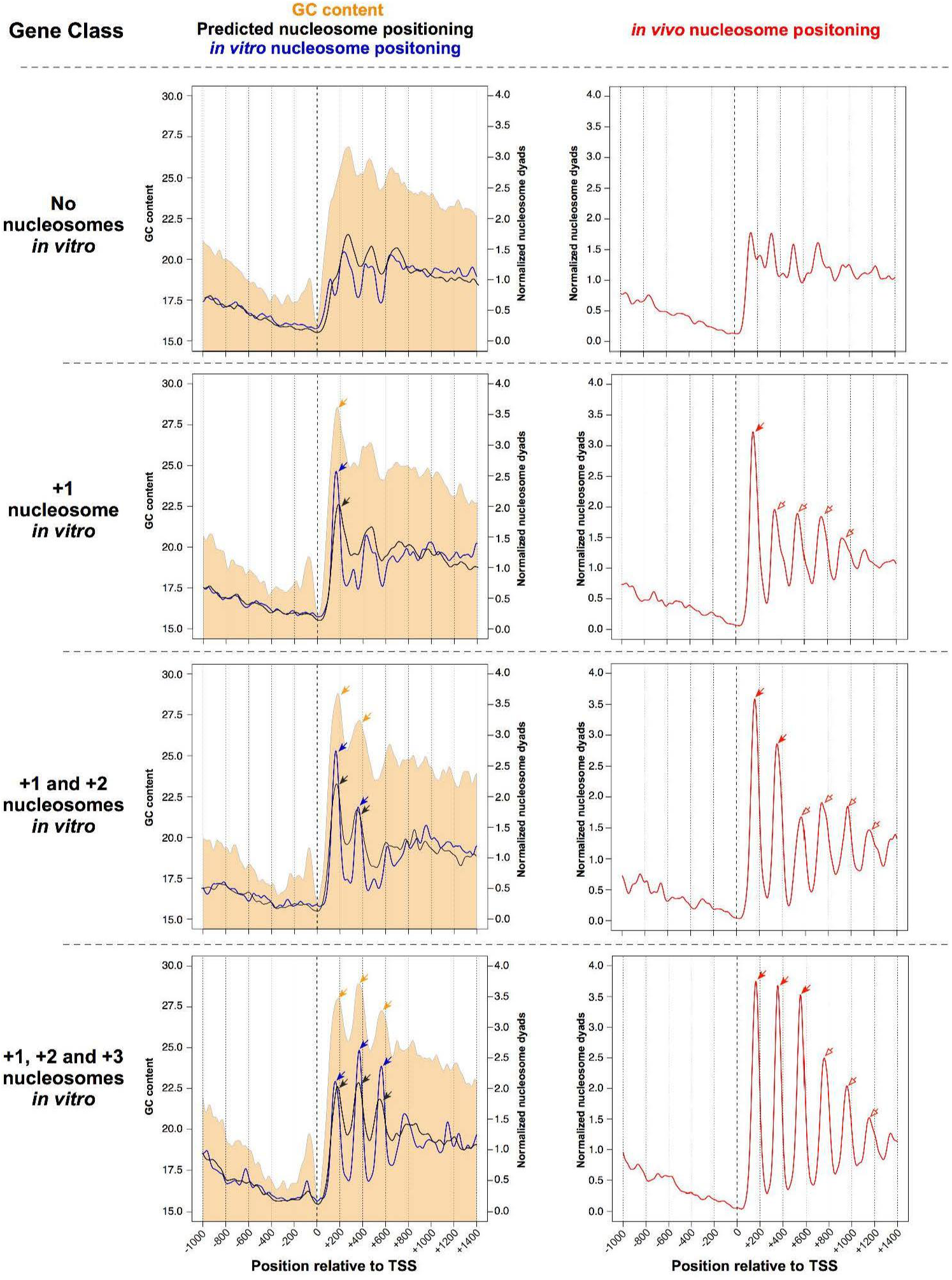
Canonical *in vitro* nucleosomes coincide with GC content oscillations, and are associated with increased nucleosome positioning *in vivo.* *Tetrahymena* genes wereclassified according to the number of canonical *in vitro* nucleosomes downstream of their TSS. Nucleosome positioning data are obtained from *in vitro* (blue line) and *in vivo* (red line) experiments, as well as from predictions of a thermodynamic model formulated by (Kaplan et al. 2009) (black line). Log-phase MNase-seq data were used as the *in vivo* sample. GC content is represented as a filled orange curve. Different gene classes are separated by horizontal dotted lines. The nucleosome-depleted region upstream of canonical nucleosomes coincides with GC-poor DNA. Pronounced peaks in GC content (orange arrows) exhibit a ~200bp periodicity, and coincide with nucleosome positions *in vitro* (blue arrows). This is consistent with GC-rich DNA being intrinsically favorable for nucleosome formation. Genes with no canonical nucleosomes *in vitro* (top row) exhibit an indistinct nucleosome pattern *in vivo* (right panel). On the other hand,genes with a +1 nucleosome *in vitro* (blue arrow within left panel) exhibit increased nucleosome positioning *in vivo*, not only at the +1 position (red filled arrow), but also around this region (red hollow arrows). A model based on nucleosome sequence preferences successfully predicts *in vitro* nucleosome positions (black arrows), which in turn overlap with *in vivo* nucleosomes (red filled arrows). However, the model fails to predict *in vivo* nucleosomes in surrounding regions (red hollow arrows), suggesting that such nucleosomes are instead positioned by *trans*-acting factors. These trends are also observed in other gene classes, with varying numbers of nucleosomes *in vitro*. DNA sequences favorable for nucleosome formation may thus function as nucleation sites that aid *trans*-acting factors in positioning nucleosomes in flanking regions *invivo*.

Next, we asked whether species-specific variation in the DNA affinity of histone octamers (Allan et al. 2013) underlies the *in vitro* pattern observed uniquely in *Tetrahymena*. We addressed this by comparing *Tetrahymena* nucleosome sequence preferences to those previously measured by *in vitro* reconstitution of chicken nucleosomes on yeast DNA. We find *in vitro* that the average nucleosome occupancies of nucleotide 5-mers correlate well between *Tetrahymena* and yeast, (Supplemental Fig. S13; Spearman ρ = 0.93). We also observe ~10bp periodic dinucleotide patterns within *Tetrahymena* nucleosomes (Supplemental Fig. S14), similar to previous analyses of yeast and human nucleosomes (Gaffney et al. 2012; Kaplan et al. 2009). Finally, we used a previously published thermodynamic model (Kaplan et al. 2009), trained on the same yeast dataset, to predict nucleosome positioning in *Tetrahymena*. We find that the genome-wide distribution of nucleosome dyads is similar between the *in vitro* dataset and predictions from the model (Spearman ρ = 0.69). These data together argue that the observed differences in nucleosome organization *in vitro* between *Tetrahymena* and yeast likely arise from distinct DNA sequence features encoded within each genome (Fig. 3), rather than species-specific DNA sequence preferences of *Tetrahymena* and yeast histone octamers. However, we cannot entirely rule out contributions from the latter possibility to the establishment of *in vivo*-like nucleosome patterns *in vitro*.

**Figure 3.**
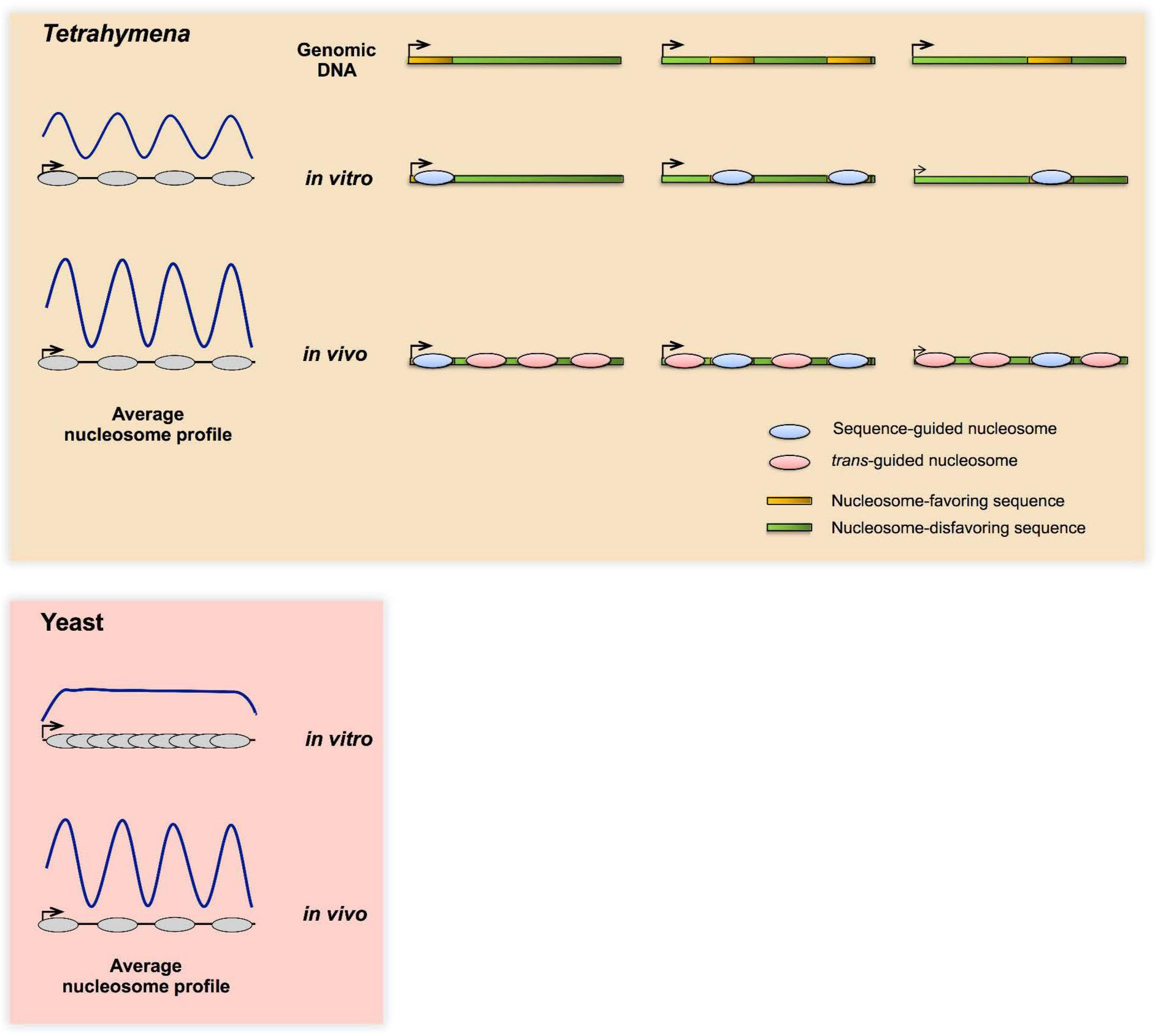
Contrasting mechanisms underlie conserved nucleosome patterns *in vivo*, between *Tetrahymena* and yeast. The *Tetrahymena* genome is GC-poor, and is generally unfavorable for nucleosome formation. The majority of *Tetrahymena* genes encode nucleosome-favoring sequences at subsets of canonical positions downstream of TSSs, which may facilitate nucleosome positioning in and around these regions *in vivo*. On the other hand, yeast genes generally show no such DNA-guided specificity near TSSs, instead relying mainly on *trans*-acting factors to generate the distinctive nucleosome organization *in vivo*. As a result, the average *in vitro* and *in vivo* nucleosome patterns appear similar in *Tetrahymena*, but not yeast.

### Sites containing DNA-guided nucleosomes exhibit greater *in vivo* positioning in their vicinity

We then addressed the *in vivo* consequences of encoding only a subset of canonical nucleosomes in the *Tetrahymena* genome. Curiously, genes with a DNA-guided nucleosome at a canonical position *in vitro* exhibited more distinct *in vivo* nucleosome positioning, at and around this location (Fig. 2). For example, genes with a +1 nucleosome *in vitro* were not only significantly more likely to possess a +1 nucleosome (p = 3.77 x 10^-14^), but also a +2 (p = 5.05 x10^-5^) and +3 nucleosome *in vivo* (p = 1.62 x 10^-3^, all with Fisher’s exact test). Similarly, those with a +2 nucleosome *in vitro* were more likely to exhibit +1 (p = 1.03 x 10^-6^), +2 (p = 1.91 x 10^-10^), and +3 nucleosomes *in vivo* (p = 6.69x10^-5^, all with Fisher’s exact test). Conversely, genes without any canonical nucleosomes *in vitro* lacked the regular pattern *in vivo* (Fig. 2). These results may suggest that DNA-guided nucleosomes – observed at canonical positions *in vitro* – act as nucleation sites to position adjacent nucleosomes *in vivo*, possibly through packing effects or the action of chromatin remodelers. Furthermore, we find that DNA-guided nucleosomes are more resistant to MNase digestion, exhibit smaller changes in translational positions between different environmental conditions, and are more strongly positioned *in vivo* than *trans* factor-guided nucleosomes (Supplemental Fig. S15A). These properties of DNA-guided nucleosomes also hold true, not only at canonical positions near TSSs, but across the entire genome (Supplemental Fig. S15B). Indeed, the genome-wide correlation in nucleosome positioning between *in vitro* and *in vivo* datasets increases with prolonged MNase digestion of chromatin, indicating that DNA-guided nucleosomes are more resistant to perturbations *in vivo* (Supplemental Fig. S16).

### DNA-guided nucleosomes impose biases on codon usage and amino acid composition

Given our findings that some canonical nucleosome positions are encoded by endogenous DNA sequences within genes, we asked how this feature is reconciled with other sequence constraints, such as the genetic code and the GC-poor nucleotide composition of the genome. Indeed, the GC content oscillations associated with DNA-guided nucleosome positioning *in vitro* overlap extensively with coding sequences, given the short 5’ UTRs of *Tetrahymena* genes (Supplemental Fig. S17). We quantified the impact of nucleosome sequence preferences on codon composition by comparing the GC content of each of the three nucleotide positions within codons that are found within DNA-guided nucleosomes, versus the corresponding nucleotide positions in codons that are found within *trans* factor-guided nucleosomes. We find that codons that overlap DNA-guided nucleosomes exhibited significantly higher GC content at all three positions (p < 2.2x10^-16^, Fisher’s exact test, for each position respectively; Table S2) than *trans* factor-guided nucleosomes. This enrichment in GC-rich codons results in deviations in amino acid composition mainly arising from the first and second codon positions, as well as deviations in synonymous codon usage from the third (wobble) position (Supplemental Tables S3 and S4). Thus, local variation in GC content – which likely underlies DNA-guided nucleosome patterns *in vivo* – imposes biases in amino acid composition and codon usage within genes.

## Discussion

In this study, we present genome-wide *in vivo* and *in vitro* nucleosome maps of the ciliate *Tetrahymena thermophila*. These maps not only constitute a comprehensive resource for further studies of ciliate chromatin (Coyne et al. 2012), but also provide novel insight into nucleosome positioning mechanisms within genes, and allude to their impact on genome evolution. The stereotypical nucleosome array that has been previously observed near transcription start sites in aggregate plots remains somewhat of a mystery. This organization has been observed in diverse eukaryotes (Yuan et al. 2005; Mavrich et al. 2008b; Lantermann et al. 2010; Ponts et al. 2010; Chen et al. 2013b; Zhang et al. 2014), suggests that it is established by a widely conserved mechanism. However, the functional relationship between the stereotypical nucleosome array and gene transcription is unclear, since even highly expressed genes exhibit this nucleosome organization (Shivaswamy et al. 2008; Lantermann et al. 2010). Paradoxically, such nucleosomes lie within coding regions near the TSS, and should thus present a significant barrier to the passage of RNA polymerase II (Teves et al. 2014). Furthermore, recent experiments in yeast demonstrated that *in vivo* ATP-dependent factors, rather than nucleosome sequence preferences, are mainly responsible for this organization. In light of these findings, it has been suggested that the stereotypical nucleosome array downstream of TSSs arises as a byproduct of a conserved process such as transcriptional elongation (Hughes et al. 2012; Struhl and Segal 2013). Our results in *Tetrahymena* now suggest that the distinct nucleosome organization may, in fact, be more than a byproduct. Our finding that some of the nucleosomes in these stereotypical arrays are guided by the underlying DNA is unexpected *per se*; yet even more striking is that they are encoded at specific stereotypical positions amidst coding sequences. Because the genetic code is highly constrained, encoding any additional information in parallel can potentially affect both codon and amino acid usage. Indeed, we demonstrate that both codon and amino acid usage are skewed at the positions where DNA-guided nucleosomes are positioned, alluding to their importance.

In previous studies, the stereotypical nucleosome array was mostly studied as a pattern averaged over many genes. This may have been a reasonable mode of analysis, since it reduces measurement noise associated with individual genes. Furthermore, individual genes in previously studied eukaryotes do indeed exhibit the array *in vivo*, consistent with the average pattern. Unexpectedly, this is not the case in *Tetrahymena*. Given our surprising observation that the stereotypical nucleosome pattern is present in the averaged pattern *in vitro*, we chose to perform further analysis at the level of individual genes. While, on average, the whole array is apparent *in vitro*, we found that individual genes mostly exhibit only subsets of these stereotypically arranged nucleosomes *in vitro*. These DNA-guided nucleosomes are more resistant to nuclease digestion, are flanked by well-positioned nucleosomes *in vivo*, and are more strongly positioned than nucleosomes guided by *trans*-acting factors. We propose that the strategic placement of these seed nucleosomes, through nucleosome-favoring sequences, could have evolved as an elegant solution to organizing nucleosomes within *Tetrahymena* genes, which are GC-poor and thus intrinsically unfavorable for nucleosome formation, while minimizing the consequences on protein-coding sequences (Fig. 3). They may thus act as nucleation sites to facilitate array formation in flanking regions *in vivo*, together with the help of *trans*-acting factors. Indeed, the notion of seed nucleosomes promoting *in vivo* nucleation of nucleosome arrays has been proposed in the human genome (Valouev et al. 2011) and could be a general mechanism for organizing chromatin within some eukaryotic genomes.

In conclusion, we find that nucleosome sequence preferences and *trans*-acting factors work together in a previously unreported fashion and extent in *Tetrahymena* to establish the distinctive nucleosome pattern in genes. These forces may function in concert with epigenetic marks such as DNA methylation, which disfavor nucleosome formation (Huff and Zilberman 2014). The arising evolutionary implications leave open the question of how distinct nucleosome positioning mechanisms operate in the context of numerous other regulatory codes enmeshed within the genome, including the maintenance of transcription factor binding sites (Stergachis et al. 2013), translational efficiency (Fredrick and Ibba 2010), mRNA splicing fidelity (Parmley et al. 2006; Parmley and Hurst 2007) and secondary structure (Shabalina et al. 2006).

## Methods

### Cell culture

1000ml of *Tetrahymena thermophila* wild-type strain SB210 (*Tetrahymena* stock center) was grown in 1xSPP at 30 °C with shaking at 100 rpm to a log-phase density of ~ 35x10^4^ cells/ml. The cell density matched that used by a recently published *Tetrahymena* RNAseq study (Xiong et al. 2012), allowing its direct integration with our MNase-seq data. To obtain starved samples, the cells were centrifuged at 1100 g for 2 min, resuspended in 1.75 volumes of 10 mM Tris pH 7.5, and incubated at 25 °C without shaking for 15 hr.

### Purification of macronuclei and MNase digestion *in vivo*

Preparations were performed as described (Jacob et al. 2004), with minor modifications. Log-phase or starved cells were centrifuged at 1000 g for 5 min, and resuspended in 70 ml TMS (10 mM Tris pH 7.5, 10 mM MgCl_2_, 3 mM CaCl_2_, 0.25 M sucrose, 1 mM DTT, 0.16 % [v/v] NP40). The culture was lysed in a Waring PBB212 Blender at the “High” setting for 25 sec, as previously described (Gorovsky et al. 1975). Formaldehyde was simultaneously added to a final concentration of 1 % (v/v), at the onset of blending. The resulting cell lysates were stirred on ice for 15 min before quenching with 125 mM glycine, and subsequent stirring on ice for an additional 10 min. These fixation and quenching steps were omitted in experiments to prepare native chromatin. Sucrose was then added at 0.816 g per ml lysate, with constant stirring on ice for 10 min. Upon complete dissolution of sucrose, the lysates were cleared by centrifugation at 9000 g for 30 min. The resulting macronuclear pellet was washed in buffer A (15 mM Tris pH 7.5, 60 mM KCl, 15 mM NaCl, 2 mM CaCl_2_, 0.5 mM spermidine trihydrochloride, 0.15 mM spermine tetrahydrochloride, 1 mM DTT), centrifuged at 1200 g for 2 min, and resuspended in 1ml buffer A. An 830 μl aliquot of macronuclei was pre-incubated at 37 °C for 5 min. From this, 100 μl macronuclei was withdrawn and mixed with 52 μl lysis buffer (300 mM NaCl, 30 mM Tris pH 8, 75 mM EDTA, 1.5 % (w/v) SDS, 1.5 mg/ml Proteinase K) to serve as an undigested control. Subsequently, 2,000 Kunitz units of MNase (NEB) were added to the macronuclei, and incubated at 37°C for 45s, 1 min 15 s, 2 min 30 s, 5 min, 7 min 30s, 10min, and 15 min respectively. At each time point, 100 μl macronuclei was withdrawn and mixed with 52 μl lysis buffer. All samples were incubated at 65 °C overnight to reverse formaldehyde crosslinks and digest proteins. DNA was subsequently purified through phenol-chloroform extraction, then ethanol-precipitated, and resuspended in buffer EB (Qiagen). 1μl of DNA from each sample was run on a 2 % agarose-TAE gel to check the progression of MNase digestion. The sample with ~80 % mononucleosomal and ~20 % dinucleosomal DNA (Supplemental Fig. S3) was labeled “light digest”. This is in accordance with previous recommendations for an adequate level of MNase digestion in nucleosome mapping studies (Zhang and Pugh 2011). Separately, the sample exhibiting almost exclusively mononucleosomal DNA with a significant smear in the subnucleosomal region (Supplemental Fig. S3) was labeled “heavy digest”. Undigested control gDNA was also sheared on a Covaris LE220. Light, heavy-digest and sheared gDNA samples were each run on a 2 % agarose-TAE gel, and the mononucleosome-sized fragment was excised and purified using a QIAquick gel extraction kit (Qiagen). Illumina libraries were prepared from mononucleosomal DNA according to manufacturer’s instructions, and subject to single-read sequencing.

### Chromatin reconstitution and MNase digestion *in vitro*

Genomic DNA for reconstitution experiments was obtained from macronuclei of starved *Tetrahymena* cells. Macronuclei were isolated from starved *Tetrahymena* cells as described earlier, and incubated in lysis buffer at 55 °C for 16 hr. Samples were purified through phenol-chloroform extraction and ethanol precipitation, and subsequently RNase-treated. ~45μg genomic DNA was then sheared to 850 bp – 2 kb using a Covaris LE220. This size range is in accordance with previously published *in vitro* reconstitution experiments (Valouev et al. 2011). Sheared DNA was end-repaired with 20 U DNA polymerase I (NEB), 60 U T4 DNA polymerase(NEB), 0.4mM dNTP, and 200 U T4 polynucleotide kinase (NEB) in a total volume of 400 μl at 20 °C for 40 min. The sample was then purified through phenol-chloroform extraction and ethanol precipitation, and then resuspended in nuclease-free water.

To obtain *Tetrahymena* histone octamers for *in vitro* reconstitution, macronuclei were first isolated from 1.4 x 10^9^ cells as described earlier. Histones were subsequently acid-extracted, as previously described (Wiley et al. 2000). Briefly, 4.39 ml 0.4 N H_2_SO_4_ was added at a ratio of 4.39 ml per 10^8^ macronuclei, and incubated at 4 °C for 3 hr with gentle shaking. The suspension was then cleared by centrifugation at 4000 g for 10 min. Perchloric acid was added to the resulting supernatant, at a final concentration of 5.4 % (w/w), and incubated on ice for 1hr. This treatment solubilized histone H1, greatly reducing contamination of the core histone preparation. Samples were then centrifuged at 4000g for 10min, and the pellet was washed with 0.1% (w/w) HCl in cold acetone, and subsequently with unacidified cold acetone. After air-drying at room temperature for 1 hr, the histone pellet was dissolved in unfolding buffer (7 M guanidinium HCl, 50 mM Tris pH 7.5, 10 mM DTT) and refolded into octamers through dialysis against 4 changes of 1L refolding buffer (2 M NaCl, 20 mM Tris pH 7.5, 1 mM EDTA pH 8, 5 mM β-mercaptoethanol) as previously described (Luger et al. 1999). Subsequently, the sample was cleared by centrifugation at 17,900 g for 5 min, before loading onto a Superdex-200 size exclusion column, equilibriated with refolding buffer. Purified histone octamer fractions were pooled, concentrated using Vivaspin 500 columns (GE Healthcare), and subsequently flash-frozen in 50 % glycerol.

Together, the sheared macronuclear genomic DNA and purified histone octamers were used for *Tetrahymena* chromatin assembly through salt gradient dialysis (Luger et al. 1999). Briefly, 3 μg of histone octamer was mixed with 7.55 μg of gDNA in a 50 μl total volume, and dialyzed against 200ml buffer C (10 mM Tris pH 7.5, 1.4 M KCl, 0.1 mM EDTA pH 7.5, 1 mM DTT) for 1 hr at 4°C. Then, 350ml buffer D (10 mM Tris pH 7.5, 10 mM KCl, 0.1 mM EDTA, 1 mM DTT) was slowly added to the assembly buffer at ~1ml/min with constant stirring. The chromatin assembly reaction was dialyzed against 200 ml fresh buffer D overnight at 4°C, followed by another change of 200 ml fresh buffer D and final dialysis for 1 hr at 4°C.

Reconstituted chromatin was adjusted to 5 mM MgCl_2_, 5 mM CaCl_2_, 70 mM KCl, and 10 mM HEPES pH 7.9 in a final volume of 60 μl. Then, 7.32 μl (22 Kunitz units) of MNase (NEB) was added to the chromatin, and incubated at 25 °C for 12 min. The digestion reaction was stopped by the addition of 21.6 μl buffer E (33 mM Tris pH 8, 100mM EDTA, 0.67 % (w/v) SDS, 16.7 % (v/v) glycerol) and 8.4 μl 20 mg/ml proteinase K. Samples were incubated at 50 °C at 1 hr. and then loaded on a 2 % agarose-TAE gel. The mononucleosome-sized fragment was gel-purified using a QIAquick gel extraction kit (Qiagen). Illumina libraries were then prepared from mononucleosomal DNA according to manufacturer’s instructions.

### MNase digestion of *Tetrahymena* gDNA

12.9 μg macronuclear gDNA was made up to 200 ul with TMS, and then digested with 1.79 Kunitz units of MNase (NEB) for 7 min at 25 °C. The reaction was terminated by adding 112 μl stop buffer (300 mM NaCl, 30 mM Tris pH8, 75 mM EDTA, 1.5 % [w/v] SDS, 1.5 mg/ml Proteinase K), and subsequently purified through phenol-chloroform extraction and ethanol precipitation. MNase-digested gDNA was resuspended in 25 μl buffer EB (Qiagen) and loaded on a 2 % agarose-TAE gel, and the mononucleosome-sized fragment was gel-purified using a QIAquick gel extraction kit (Qiagen). Illumina libraries were then prepared from gDNA according to manufacturer’s instructions.

### Sequencing data processing pipeline

All Illumina sequencing datasets are summarized in Supplementary Table S5. Raw MNase-seq and genomic DNA-seq reads were quality-trimmed (minimum quality score = 20) and length-filtered (minimum length = 40nt) using Galaxy (Giardine et al. 2005; Blankenberg et al. 2010; Goecks et al. 2010) before mapping with BWA (Li and Durbin 2009) to the October2008 build of the *Tetrahymena* SB210 reference genome (Eisen et al. 2006) using standard settings. Only complete *Tetrahymena* chromosomes in the genome assembly were included in downstream analyses. We then used the sheared genomic DNA-seq data to calculate RPKM values (reads per kb per million mapped reads) for each chromosome in log-phase and starved conditions, respectively. These RPKM values were used as a measure of relative copy number of *Tetrahymena* chromosomes, which are highly polyploidy (Eisen et al. 2006) (~45n). In addition, because nuclear division proceeds in the absence of a mitotic spindle (Lauth et al. 1976), inter-chromosomal variation in copy number may exist. This could affect calculations of nucleosome positioning between different chromosomes. Relative DNA copy number data were subsequently used to normalize MNase-seq data, as described next.

Average fragment sizes for each MNase-seq dataset (including MNase-digested naked DNA) were calculated using cross-correlation analysis (Kharchenko et al. 2008) (see Supplemental Table S2). Reads in each dataset were then extended in length to match their respective inferred fragment sizes. The center of each extended read was designated as the nucleosome dyad position. Per-basepair coverage of nucleosome dyads was calculated across the *Tetrahymena* genome for each dataset. Following this, the data were normalized by relative chromosome copy number (as obtained from RPKM of sheared genomic DNA-seq reads for each chromosome), and the whole genome average coverage value. Normalized values were then smoothed with a Gaussian filter of standard deviation = 15. We refer to the resulting values as normalized nucleosome dyads.

RNAseq data from log-phase and 15 hr-starved *Tetrahymena* was obtained from a published study (Xiong et al. 2012). Reads were first quality-trimmed and length-filtered using cutadapt (parameters: “-e 0.1 -O 8 -m 25 -q 20”). Subsequently, they were mapped to Tetrahymena rDNA and mitochondrial sequences using BLAT (parameters: “-noHead - stepSize=5 -minIdentity=92”). RNAseq reads not of rDNA or mitochondrial origin were then mapped to *Tetrahymena* genes using BLAT with the same parameters. Gene annotations: Feb2014, Genbank AAGF03000000 (Bidwell et al.) were obtained as a pre-release version from R. Coyne. DESeq was then used to calculate size factors, in order to account for differences in sequencing depth between log-phase and starve RNAseq libraries (size factors = 1.0983709, 0.6207299 respectively). Mapped read counts in each dataset were normalized by gene length and DESeq size factors to obtain relative expression values.

### Nucleosomal dinucleotide frequencies

AA/TT/TA/AT dinucleotide frequencies within nucleosomal DNA were calculated as previously described (Kaplan et al. 2009). Briefly, extended MNase-seq reads were reverse complemented, and – together with the original reads – aligned according to their start position. At each position *i* in the alignment, we calculated the average frequency of AA/AT/TA/AT dinucleotides at the [*i-1*, *i*, *i+1*] positions, representing a smoothed 3bp-sliding window. The calculated dinucleotide frequency at position *i* was subsequently normalized to the average dinucleotide frequency across all positions along the nucleosome, and then divided by the frequency of all dinucleotides at that position.

### Phasograms

We calculated the per-basepair number of read start positions across the genome, for *invivo* and *in vitro* MNase-seq data, respectively. These data were normalized by the genome-wide average number of read start positions, and smoothed with a Gaussian filter of standard deviation = 15. We then extracted the normalized per-basepair read start counts 1000bp upstream of each read start position. These data were averaged across all read starts, and plotted as the phasogram.

### Nucleosome calling

We employed a previously published iterative search procedure (Kaplan et al. 2010b)with minor modifications to identify nucleosomes based on normalized nucleosome dyad data. Briefly, the position with highest normalized read center coverage was identified, and designated as a nucleosome dyad. The flanking 140bp (for *Tetrahymena*) or 106bp (for *S.cerevisiae*) from either direction of the nucleosome dyad was then excluded to account for the nucleosome width and linker region. A smaller exclusion distance was instated for *S. cerevisiae*, given its shorter nucleosome repeat length (Lantermann et al. 2010). This process was repeated until no new global maxima were found. Called nucleosomes (peaks) were then filtered according to two published metrics (Kaplan et al. 2010a): absolute nucleosome positioning and conditional nucleosome positioning. Absolute nucleosome positioning was defined as the number of MNase-seq read centers (normalized by chromosome copy number and the genome-wide average value) that correspond to a particular peak. Conditional nucleosome positioning was defined as the normalized number of read centers that lie within 21 bp of the called nucleosome peak, divided by the normalized number of read centers that lie within 147 bp from the peak. To construct the histograms in Fig. 1 and Supplemental Fig. S9B, approximately 35 % of originally called nucleosomes were first removed using a stringent filter of minimum absolute positioning and conditional positioning. For all other analyses and Figures, nucleosomes with absolute positioning < 0.19 and conditional positioning < 0.23 were first omitted, resulting in the removal of ~15 % of peaks.

### Nucleosome model

To predict nucleosome positioning from DNA sequence, we used the thermodynamic model of (Kaplan et al. 2009), which was trained on MNase-seq data measured on chicken histones that were reconstituted onto yeast DNA. We used the same concentration and temperature parameters as used previously. In order to produce a track which is comparable to our smoothed dyad track, we applied a Gaussian filter (standard deviation = 15bp) to the track of nucleosome start probabilities given by the model and shifted it by 73 bp.

### Data access

All sequencing data generated for this study have been deposited in the NCBI Gene Expression Omnibus (GEO) under accession ID GSE64061. Previously published *Tetrahymena thermophila* RNAseq data (Xiong et al. 2012) were retrieved from the NCBI GEO under accession ID GSE27971. All yeast MNase-seq data used in this work were obtained from a previous study by Kaplan et al. 2009, under accession ID GSE13622.

## Acknowledgments

This paper is dedicated to the late Jonathan Widom, whose work inspired L. Y. B. and N. K. We thank Job Dekker, Jason Lieb, Richard Bártfai, Nicole Francis, Galia Debelouchina, Jaspreet Khurana and Geoff Dann for feedback and comments on the manuscript; Robert Coyne for sharing *Tetrahymena* gene annotations; David Robinson for advice on MNase-seq read subsampling; Jingmei Wang for general laboratory support; Jessica Wiggins, Wei Wang, and Donna Storton for assistance with Illumina sequencing. This study was supported by a Princeton Centennial Fellowship to L. Y. B., Human Frontier Science Program Long Term Fellowship LT000706/2012 to N. K. and NIH grant GM59708 to L. F. L.

## Author contributions

L. Y. B. conceived the project and performed experimental and bioinformatic analysis for all Figures and Tables. N. K. provided advice, analysis and critical interpretations of bioinformatic data. M. M. M. and T. W. M. contributed feedback, core protocols, and equipment for *in vitro* reconstitution experiments. L. F. L provided guidance and feedback. L. Y. B., and N. K. prepared the manuscript, which all authors edited.

## Supplemental Material

**Figure S1. Workflow of *Tetrahymena* nucleosome mapping experiments.** Macronuclei were isolated from starved or log-phase *Tetrahymena* and digested with MNase. Separately, *Tetrahymena* histones were acid-extracted, refolded into octamers, assembled on genomicDNA through salt gradient dialysis, and subsequently treated with MNase. No *trans*-acting factors were added during chromatin assembly. The mononucleosomal DNA from *in vivo* and *in vitro* MNase digests was gel-purified for subsequent Illumina sequencing.

**Figure S2. Subsampling of MNase-seq data.** Varying fractions of mapped reads from each dataset were randomly subsampled, and used for nucleosome calling through DANPOS (Chen et al. 2013a). Reads were mapped to all chromosomes in the October 2008 build of the *Tetrahymena* SB210 reference genome (Eisen et al. 2006), including those not capped with telomeres. The number of called high confidence nucleosomes (p <1e^-5^) approached saturation before full sampling of *in vivo* and *in vitro* data, indicating that nucleosomes are well-sampled in all datasets.

**Figure S3. Gel analysis of *Tetrahymena* chromatin.** (A) Macronuclei from log-phase or starved cells yielded nucleosome ladders upon MNase digestion *in vivo*, similar to other eukaryotes. A protected mononucleosome-sized fragment was observed after *in vitro* reconstituted chromatin after MNase treatment, with no evidence of laddering. Mononucleosomal DNA samples marked with a red arrow were gel-purified for subsequent Illumina sequencing. (B) Size exclusion chromatography of refolded *Tetrahymena* histone octamers. The fractions highlighted with a horizontal black bar were pooled and concentrated for subsequent *in vitro* reconstitution experiments with *Tetrahymena* genomic DNA.

**Figure S4. Nucleosome dyad counts along the 5’ NTS of the *Tetrahymena* ribosomal DNA locus.** Only uniquely-mapping reads were considered when tabulating nucleosome dyads from MNase-seq reads, at this locus. Blue and green tracks represent *in vivo* data from fixed or native chromatin, digested to different extents with MNase. Well-positioned nucleosomes *in vivo* flank both origins of replication *in vivo*, corroborating independent studies that mapped nucleosome positions through Southern analysis.

**Figure S5. Nucleosome organization near TSSs is similar *in vitro* and *in vivo*.** Averaged nucleosome dyad counts around the TSS (top panel) reveal an *in vivo*-like distribution of called nucleosomes within *in vitro* data. MNase-digested naked DNA does not resemble *in vivo* data (green curve), thus ruling out potential sequence biases associated with MNase preferences.

**Figure S6. Nucleosome density downstream of TSSs varies with gene expression.** *Tetrahymena* genes were binned in quintiles, based on normalized RNAseq expression in the log-phase and starved conditions. Highly expressed genes tend to exhibit increased nucleosome density downstream of TSSs, in the log-phase nutritional condition. This trend is less apparent in starved cells.

**Figure S7. Nucleosome density in promoters is negatively correlated with gene expression.** Promoter nucleosome density is calculated as the average number of nucleosome dyads between -250bp to 0bp from TSSs. Genes were binned into quintiles, based on normalized RNAseq expression in either the log-phase or starve condition. Genes with low expression levels tend to have higher nucleosome density in their promoter regions, though this trend is also less apparent in the starved nutritional condition.

**Figure S8. Removing duplicate reads does not affect the in *vivo*-like nucleosome organization near TSSs *in vitro*.** (A) Nucleosome count data were computed from the original datasets (red curve), and when duplicate reads are removed (blue curve). (B) Histograms of called nucleosome dyads within in vitro and log-phase in vivo datasets, around the TSS. Stringent filters for absolute and conditional nucleosome positioning were applied, such that~35% of nucleosomes were discarded. The nucleosome organization in both (A) and (B) remain qualitatively similar *in vivo* and *in vitro* even when duplicate reads are removed, suggesting that it is not an artifact arising from over-amplification of Illumina libraries.

**Figure S9. The phased pattern of *in vitro* nucleosome positions is robust to variation in nucleosome calling criteria.** Cutoffs for absolute positioning (abs. pos.) and conditional positioning (cond. pos.) were separately varied, such that up to 30% of called nucleosomes were respectively removed. The filtered data were then used to plot histograms of called nucleosome positions, relative to the TSS.

**Figure S10. Sites closest to the TSS show greatest correspondence between *in vitro* and *in vivo* nucleosome positions.** For *in vitro* nucleosomes in the +1, +2, +3, and +4 positions downstream of the TSS, the distance to the nearest *in vivo* nucleosome is calculated. “Other” represents *in vitro* nucleosomes not located at +1 to +4 positions. Nucleosomes at the +1 position *in vitro* most closely overlap with a nucleosome *in vivo*, suggesting that the +1 nucleosome is most greatly stabilized by DNA sequences.

**Figure S11. Phasograms of *in vitro* and *in vivo* MNase-seq datasets.** A distinct 200bp periodicity is specifically observed within *in vivo* datasets (log-phase and starve), suggesting the presence of regular nucleosome arrays. This is consistent with our gel analysis (Supplemental Fig. S3A) and other independent studies (Gorovsky et al. 1978).

**Figure S12. The *in vitro* nucleosome organization at individual genes resembles *in vivo* patterns.** Vertical black arrows represent the TSS, while light purple boxes represent the 5’UTR. Black lines indicate the presence of *in vitro* nucleosomes at *in vivo*-like locations. A minority of genes (eg. *mRNA0064.207*) exhibit *in vitro* nucleosomes at all canonical positions, while most genes have such nucleosomes at only a subset of positions.

**Figure S13. Comparison of normalized nucleosome occupancy of 5-mers in the yeast and *Tetrahymena* genomes.** Occupancy data were calculated from the number of extended *in vitro* MNase-seq reads that span each unique 5-mer, normalized by the average 5-mer read count within each genome. This represents the relative intrinsic affinities of histone octamers for various unique DNA sequences. A strong correlation between *Tetrahymena* and yeast nucleosome occupancies is observed, indicating that histone octamers from both species share similar nucleosome sequence preferences. Colored data points progressing from dark blue to red denote increasing AT content.

**Figure S14. Rotational positioning of *Tetrahymena* nucleosomes.** AA/TT/AT/TA dinucleotide frequencies were calculated as a 3bp sliding window average across nucleosomal DNA. A clear 10bp periodicity is observed, and is more distinct *in vitro* than *in vivo*. This is consistent with the larger role that nucleosome sequence preferences play in guiding nucleosome positions *in vitro*.

**Figure S15. DNA-guided nucleosomes are more resistant to nuclease digestion, exhibit less variability in translational positions between different nutritional conditions, and are more strongly positioned *in vivo*.** A nucleosome *in vivo* is classified as “DNA-guided” if the it lies within 10 bp from nucleosome *in vitro*. On the other hand, a nucleosome *in vivo* that lies greater than 73 bp from a nucleosome *in vitro* is classified as “*trans* factor-guided”. Nuclease resistance was calculated as the total number of mid-points of MNase-seq reads that lie within 73 bp of a nucleosome in heavily digested chromatin, divided by the corresponding number of MNase-seq read mid-points in lightly digested chromatin (see Methods for description of ‘heavy’ and ‘light’ chromatin digests). For every *in vivo* nucleosome in a particular environmental condition (eg. log-phase), its distance to the nearest nucleosome in another environmental condition (eg. starve) is calculated. These distances are tabulated for all DNA-guided and *trans* factor-guided nucleosomes, respectively, and are denoted as the variability in positioning between different environmental conditions. Absolute nucleosome positioning and conditional nucleosome positioning are calculated as described in Methods. (A) Analysis specifically of canonical +1, +2, and +3 nucleosomes downstream of the TSS. (B) Analysis of all nucleosomes across the genome in log-phase and starve conditions, respectively.

**Figure S16. Genome-wide comparisons of *Tetrahymena* nucleosome organization.** Pairwise whole-genome spearman correlation between various nucleosome maps. Each nucleosome map is represented as a series of nucleosome dyad counts for each basepair in the genome, normalized by the genome-wide average number of nucleosome dyads, and subsequently smoothed with a Gaussian filter of standard deviation = 15. This pipeline was also applied to the MNase-digested naked DNA sample, to maintain consistency in data processing. The correlation between *in vivo* and *in vitro* data improves with increased MNase digestion of chromatin *in vivo*. This is observed in both log-phase and starved conditions.

**Figure S17. Distribution of *Tetrahymena* ORF start positions.** Bars shaded in blue represent ORF start positions that lie upstream of the +1 nucleosome dyad, while red bars represent ORF start positions downstream of it. Most 5’ UTRs (given by the distance between the TSS and the ORF start position) are short, with a median length of 84 bp. Thus, nucleosomes downstream of the TSS likely lie within *Tetrahymena* open reading frames.

**Table S1. Numbers of genes exhibiting at least 1, 2, or 3 canonical nucleosomes at canonical positions immediately downstream of the TSS.** Respective nucleosomes were scored if they were ≤35bp from the canonical position. The *in vivo* (log-phase) +1, +2, and +3 positions are +150, +342, and +541 respectively, while the *in vitro* positions are +163, +363, and +558 respectively. A total of 2413 genes were analyzed. The absolute number of classified genes for *in vivo* and *in vitro* conditions is respectively shown. Percentages denote the fraction of genes with canonical nucleosomes *in vitro* and *in vivo*, relative to the total number of genes that have canonical nucleosomes *in vivo*.

**Table S2. Codons within DNA-guided nucleosomes exhibit higher GC content than those within *trans* factor-guided nucleosomes.** Both types of nucleosomes are defined in Supplemental Fig. S15. Codons that lie no greater than 73 bp from a called nucleosome are considered as lying within the corresponding DNA-guided or *trans* factor-guided nucleosome.

**Table S3. Biases in synonymous codon usage are encoded within distinct nucleosomal regions.** DNA-guided and *trans*-factor guided nucleosomes are defined as in Supplemental Fig. S15. Codons were considered as lying within a nucleosome, according to criteria described in Supplemental Table S2. Each group of synonymous codons was analyzed separately. Codons with high GC content relative to synonymous counterparts are shaded red, while those with low GC content are shaded blue. Separately, codons enriched within DNA-guided nucleosomes are highlighted in red, while those depleted are highlighted in blue. This enrichment/depletion value was calculated by dividing the codon frequency in sequences that lie within 10 bp of DNA-guided nucleosomes, by the codon frequency within sequences that lie within 10 bp of *trans* factor-guided nucleosomes. It quantifies the impact of accommodating DNA-guided nucleosomes on synonymous codon usage. The underlying codon usage for 15 out of 18 amino acids was biased towards GC-rich codons within coding regions that overlap with DNA-guided nucleosomes.

**Table S4. Biases in amino acid composition are encoded within distinct nucleosomal regions.** DNA-guided and *trans*-factor guided nucleosomes are defined as in Supplemental Fig. S15. Amino acids whose corresponding codons lie no greater than 73 bp from a called nucleosome are considered as lying within the nucleosome. Weighted codon GC content values were calculated as the sum of GC contents of synonymous codons specifying an amino acid, respectively normalized by their respective codon frequencies. Amino acids were ranked according to their weighted codon GC content, as shaded from low (blue) to high (red). Amino acids specified by GC-rich codons tend to be enriched in coding regions that overlap with DNA-guided nucleosomes.

**Table S5. Average fragment sizes and sequencing read depth of Illumina datasets used in this study.** Fragment size data were calculated using cross-correlation analysis (Kharchenko et al. 2008). Sequencing reads counts denote the total number of reads mapped to two-telomere (complete) chromosomes in the *Tetrahymena* SB210 genome assembly.

